# Maternal effects influence temperature-dependent offspring survival in *Drosophila melanogaster*

**DOI:** 10.1101/372870

**Authors:** Snigdha Mohan, Ton G.G. Groothuis, Chris Vinke, Jean-Christophe Billeter

## Abstract

Mothers may modulate the phenotype of their offspring by affecting their development based on her own environment. In changing environments, these maternal effects are thought to adjust offspring physiology and development and thus produce offspring better prepared to the environment experienced by the mother. However, evidence for this is scarce. Here we test the consequences of a match or mismatch between mother and offspring temperature conditions on growth, adult morphology and reproduction into the grandchildren generation in the fruit fl*y Drosophila melanogaster*. This experimental design tests the relative contribution of maternal effects and offspring intrinsic plasticity to the phenotypic response to temperature conditions. We manipulated maternal temperature conditions by exposing mothers to either 18°C or 29°C conditions. Their eggs developed at a temperature that was either matched or mismatched with the maternal one. Survival from egg to adult was higher when the maternal and offspring environments matched, showing maternal effects affecting a trait that is a close proxy for fitness. However developmental speed, adult size and fecundity responded to temperature mostly through offspring phenotypic plasticity and maternal effects only had a small contribution. The results provide experimental evidence for maternal effects in influencing a potentially adaptive offspring response to temperature in the model organism *Drosophila melanogaster.* These effects appear to modulate early embryonic phenotypes such as survival, more than the adult phenotypes of the offspring.

## Introduction

Changes in biotic and abiotic conditions are a normal feature of most environments. Organisms can adjust to these changes through genetic variants, or, in the time frame of one lifetime. through developmental, physiological and behavioral phenotypic plasticity. This plasticity allows the emergence of different phenotypes and life history strategies adapted to specific environmental variables (Nylin, 2013). Phenotypic plasticity often arises through mechanisms modulating developmental events. As these mechanisms occur during early development, embryos might not yet be equipped to sense environmental cues predicting its later environment. A route for controlling phenotypic plasticity is via the parents, who experience cues of environmental change and may adjust offspring development by influencing their prenatal environment. This can be achieved by influencing egg composition or the transfer of nutrients, immune factors or hormonal signals during pregnancy that can induce epigenetic changes regulating developmental plasticity and resulting in phenotypic differences in the offspring (Groothuis *et al.*, 2005).

An outstanding question is to what extend phenotypic plasticity is based on cues experienced by the individual versus cues experienced by their parents (Uller *et al.*, 2013; Groothuis & Taborsky, 2015). If plasticity in a particular phenotype is adaptive and can be traced back to parental effects, induced by the parental environment, then this indicates that the parents have made adjustments relevant to the postnatal environment of their offspring. In this case the parental prediction of the offspring environment is then accurate, the offspring’s phenotype will “match” the environment in which it will live, potentially increasing its fitness. However, if the prediction is wrong, there is a ‘mismatch’ at the potential cost of the survival and/or fecundity of the offspring. However, environmental conditions can also directly affect the parents ability to provision their eggs, or look after their offspring. Such effects can carry over to their offspring but do not represent anticipatory plasticity as the parental experience, such as food and resource limitation, simply carry over to the next generation and constrain their development (Uller *et al.*, 2013; Nettle & Bateson, 2015; Raveh *et al.*, 2016; Engqvist & Reinhold, 2016). There are few clear examples of anticipatory parental effect. For instance, in daphnia, parents exposed to predators produce offspring that are morphologically better equipped against predation (Agrawal *et al.*, 1999). The broad adaptive relevance of such anticipatory parental effects however remains controversial, in part because of the methodological difficulties in finding the right environmental cues and the requirement of testing phenotypes of offspring in full factorial design including exposing and testing offspring in environments that are either matched or mismatched with that of their parents (Uller *et al.*, 2013).

We developed a paradigm to test the anticipatory nature of parental effects in laboratory conditions, allowing measuring the separate contribution of parental effect and direct environmental effects on offspring phenotypic plasticity. We chose to study the effect of ambient temperature because it is an environmental variable that fulfills three criteria for testing anticipatory parental effects: it is not constant, its changes are related to seasons and thus predictable for the mother, and it is sufficiently persistent to be of relevance for developmental phenotypic adjustment. Moreover, temperature induces transgenerational effects in several species, including fish (Salinas & Munch, 2012; Munday, 2014). We chose to study the fruit fly *Drosophila melanogaster* because its development is strongly temperature dependent, its cosmopolitan distribution exposes it to a large range of temperatures and substantial fluctuations in temperature over the reproductive season depending on its geographical location (Hoffmann, 2010). The fast generation time of this species (7 days at 29°C; (Ashburner, 1989) means that environmental variables experienced by parents may match those of the postnatal environment of their offspring, making anticipatory maternal effects a potentially relevant mechanism.

*Drosophila* has behavioural and morphological phenotypic plasticity in response to temperature (James *et al.*, 1997; Gilchrist & Huey, 2001; Petavy *et al.*, 2001; Trotta *et al.*, 2006). For example, flies developing at 18°C will develop slower but reach larger adult size than genetically identical flies developing at 29°C. However, flies housed in hotter conditions are typically more fecund than those in colder conditions (Kingsolver & Huey, 2008). Previous studies have described parental effects in *Drosophila* linked to temperature on a variety of traits including developmental speed (Huey *et al.*, 1995; Gilchrist & Huey, 2001), cold tolerance (Watson & Hoffmann, 1995), egg size (Crill *et al.*, 1996) and survival (Magiafoglou & Hoffmann, 2003), but the one study that tested parental effects in a match-mismatch design did not find evidence that a match between parent and offspring environment resulted in greater offspring fitness. However, developmental survival, an important proxy for fitness, was not measured in those studies.

Here we tested the relative contribution of parental effected and offspring phenotypic plasticity in *Drosophila* up to the second generation using a full factorial match-mismatch design. We exposed mothers to one of two temperature conditions (18°C and 29°C) and let their offspring develop under either matched or mismatched temperatures (Figure 1). We examined the effect of match-mismatch conditions on offspring morphological traits (such as egg size\volume and wing size), and life history traits (such as survival, fecundity and developmental time), to estimate the size of parental effects and offspring intrinsic phenotypic plasticity of these traits in different stages of development.

**Figure 1:**
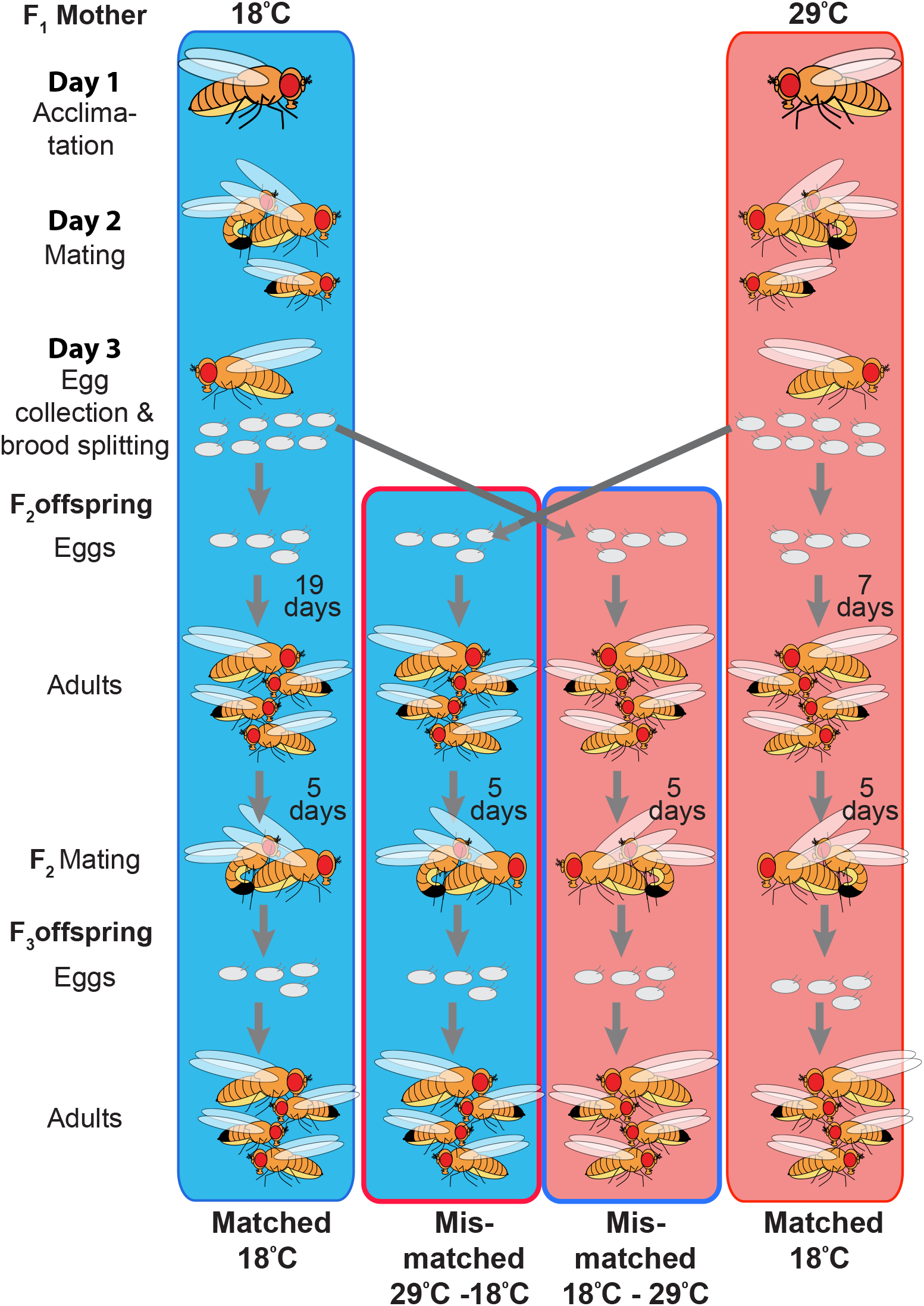
Match-Mismatch design to investigate anticipatory parental effects in response to temperature conditions. Newly emerged F_1_ adult females who developed at 25°C were acclimated to 18°C or 29°C for 24 hours. Females were then housed for 24 hours with two males for fertilization. Males were discarded and the females were allowed to lay eggs for 24 hours. Eggs were collected and split in four groups: Matched 18°C group, where mothers experienced 18°C condition and offspring developed at 18°C; Mismatched 29°C-18°C group, where mothers experienced 29°C and offspring developed at 18°C; Matched 29°C group, where mothers experienced 29°C and offspring developed at 29°C; Mismatched 18°C −29°C group, where mothers experienced 18°C and offspring developed at 29°C. Eggs were transferred to a food vial where they developed until adulthood. Arrows from F_2_ eggs to adults indicates the developmental time in matched conditions. Pairs of F_2_ adult males and females were mated at the same temperature they developed. Their F_3_ offspring were also raised at those same temperatures.

## Material and Methods

### *Drosophila* stocks and rearing conditions

The *Oregon-R* laboratory wild-type strain was used in all experiments. Stocks were kept in vials at 25°C in a 12:12 Light-Dark (LD) cycle and reared on fly food (referred henceforth as “food”) medium containing agar (10g/L), glucose (167mM), sucrose (44mM), yeast (35g/L), cornmeal (15g/L), wheat germ (10g/L), soya flour (10 g/L), molasses (30 g/L), propionic acid and Tegosept. For Temperature treatment, flies were reared in two walk-in climate chambers, one set at 18°C (average recorded temperature 17.7°C, with min at 17.3 and max at 18.3) and one set at 29°C (average recorded temperature 28.7°C, with min of 28.3°C and max of 29.8°C).

### Experimental design: Match-Mismatch temperature treatment

#### Generation of F_1_

The experimental treatments schedule is outlined in fig. 1. Approximately 200 F_0_ flies were placed in an egg-laying cage with a removable egg-laying dish. The egg laying dish consisted of a 35×10mm petri dish layered with 3 ml of a solution composed of 20g agar, 26g sucrose, 52g glucose, and 9% (v/v) red grape juice per litre of distilled water spotted with a fresh dab of dry yeast mixed with water. The cage was kept at 25°C in a 12:12LD incubator. Eggs were collected twice a day at Circadian Time (CT) 0 and CT8 by replacing the egg-laying dish. Larvae were picked 24hr later from dishes stored at 25°C. Groups of 40 larvae were transferred to a single 25×95mm plastic vial containing 6ml of food (referred to as food vial) and left to develop to adulthood at 25°C in a 12:12LD incubator. Virgin F_1_ females were collected from these vials at room temperature (~22°C) using mild CO_2_ anesthesia (exposure for maximum couple of minutes under minimal CO_2_ flow).

#### Treatment of F_1_

F_1_ virgin females were individually transferred immediately after collection to a 35×10mm Petri dish layered with 3 ml of food. The dishes were moved within an hour of collection to either an 18°C or to a 29°C walk-in climate chamber with a 12:12LD cycle. After 24 hours, two virgin males, offspring of the same F_0_ flies, that had been raised and aged at 25°C in a 12:12LD incubator, were added to each dish to fertilize the females. Twenty-four hours later, single females were transferred to individual dishes with fresh fly food and a dab of yeast paste to stimulate egg laying. Females were then allowed to lay eggs for 24hrs in either 18°C or 29°C conditions.

#### Treatment of F_2_

Eggs laid by F_1_ females at 18°C or 29°C were collected directly from the egg-laying dish on this third treatment day and transferred to a vial containing 6.5 ml of food for development. The brood was split by transferring half the eggs to the 18°C treatment and the other half to the 29°C treatment (fig. 1). F_2_ adults were collected at eclosion. Mating assays were performed at the same temperature at which the offspring developed and were set up by introducing one virgin female with one virgin male into a Petri dish layered with food. F_2_ siblings treated in either matched or mismatched conditions were mated with each other. After a single mating, females were transferred to food vials housed at the same temperature at which they developed to lay eggs. Females were transferred three times to a fresh vial at two days intervals to prevent overcrowding of the food vials by larvae. The number of F_3_ adults was counted at eclosion.

### Offspring traits

#### Number of eggs

The number of eggs laid at 18°C and 29°C during a 24hr egg-laying period was counted directly in the egg-laying dish.

#### Egg volume measurement

One to five freshly laid eggs were collected hourly from single females at both 18°C and 29°C (from 11 and 31 females respectively). The size of matched eggs was measured immediately at collection. To rule out a potential direct effect of temperature on egg size shortly after laying, mismatched eggs were measured 5 hrs after collection, to allow time for temperature to potentially impact egg volume, and compared to matched eggs. Eggs were photographed using a Leica MZ10F stereomicroscope equipped with a Leica DFC450c camera connected to a computer running the Leica Application Suit software. Egg Length (L) and width (W) were determined using the software ImageJ (National Institutes of Health, Bethesda, MD, USA) on photographs taken at 6.3X magnification. The volume (V) was determined by using formula V=(1/6)πW^2^L (Markow *et al*., 2009).

#### Survival from egg to adult

Eggs were collected as described above from single females at 18°C or 29°C, except that the egg collection was limited to a single 4-hour interval. Slow egg laying by females at 18°C resulted in an average of 7.5 (± 4.7) eggs collected per female (n= 81), while faster egg laying at 29°C resulted in 37.6 (± 20.4) per female (n=81). Because of the small number of eggs in this specific experimental setup, broods from single females were not split, but instead randomly assigned to 18°C or 29°C conditions after transfer to a food vial. Number of adults produced from these eggs was counted at eclosion to determine the percent survival from egg to adult.

#### Developmental Time

To determine the developmental time from egg to adult, the time and date of laying of eggs and that of adult eclosion were recorded. Groups of 15-40 eggs per female were collected at 8-16 hours interval and transferred to a food vial. This time interval was required to collect sufficient amount of eggs at 18°C, where egg-laying rate is slower than at 29°C (Huey *et al.*, 1995). Development time was determined from the time eggs were collected to the time the last adult from that group of eggs emerged.

To determine developmental time at 29°C more precisely, as development is faster under this condition than at 18°C, single eggs were collected at one hour intervals and exposed to matched or mismatched treatments. At the pupal stage, a Logitech webcam controlled by the SecurityMonitor Pro software took pictures at 1-hour intervals to determine the precise eclosion time. Red light was utilized to visualize pupae during the dark phase. These data were used to confirm developmental time differences in 29°C Match and 18°C −29°C mismatched conditions.

#### Wing size measurement

There is an association between size, fecundity and mating success in *Drosophila*; larger individuals have more offspring and have a greater chance of mating (Kingsolver & Huey, 2008). We estimated the size of matched and mismatched adult offspring as an indirect measurement of fitness. We measured wing size parameters since those are correlated with total body size and can be more accurately measured. The right wing of 5 F_2_ adults from the same mother were measured to constitute one replicate. Wings were removed with fine forceps 5-6 hours post-eclosion and mounted on a glass slide with a cover slip. Pictures of wings were taken as for egg volume. Measurement method was adapted from (Joubert & Bijlsma, 2010). Wing length and width were measured with the program ImageJ (v. 6.4).

#### Reproductive performance

The fitness of F_1_ mothers was estimated based on the number of grand-children they obtained when their offspring had been kept in conditions that matched or mismatched theirs. Three pairs of matched and three pairs of mismatched F_2_ males and females per F_1_ mother were allowed to mate a single time after which single F_2_ mated females were transferred to a fresh food vial and allowed to lay eggs for their entire lifespan. The resulting F_3_ adults were counted to determine the F_2_ reproductive performance. F_2_ and F_3_ individuals were continuously kept in the same conditions in which the original F_2_ eggs were treated, leading to an unbroken chain of matched or mismatched conditions with respect to the F_1_ maternal condition. F_2_ flies were kept at the same temperature condition in which they developed in food vials in groups of 10 individuals of the same sex for 5 days before mating.

As we did not measure lifetime reproductive output of F_1_ mothers, we used the number of F_2_ adults generated by 1 day of F_1_ mothers egg laying to estimate their reproductive output when their offspring are in matched *vs.* mismatched conditions (fig. 2*c*). The number of F_3_ produced by F_2_ was determined as described in the paragraph above. The average number of offspring for a single F_1_ mother was multiplied by the average number of offspring of single F_2_ mothers to determine reproductive performance in different temperatures and in matched or mismatched conditions.

**Figure 2:**
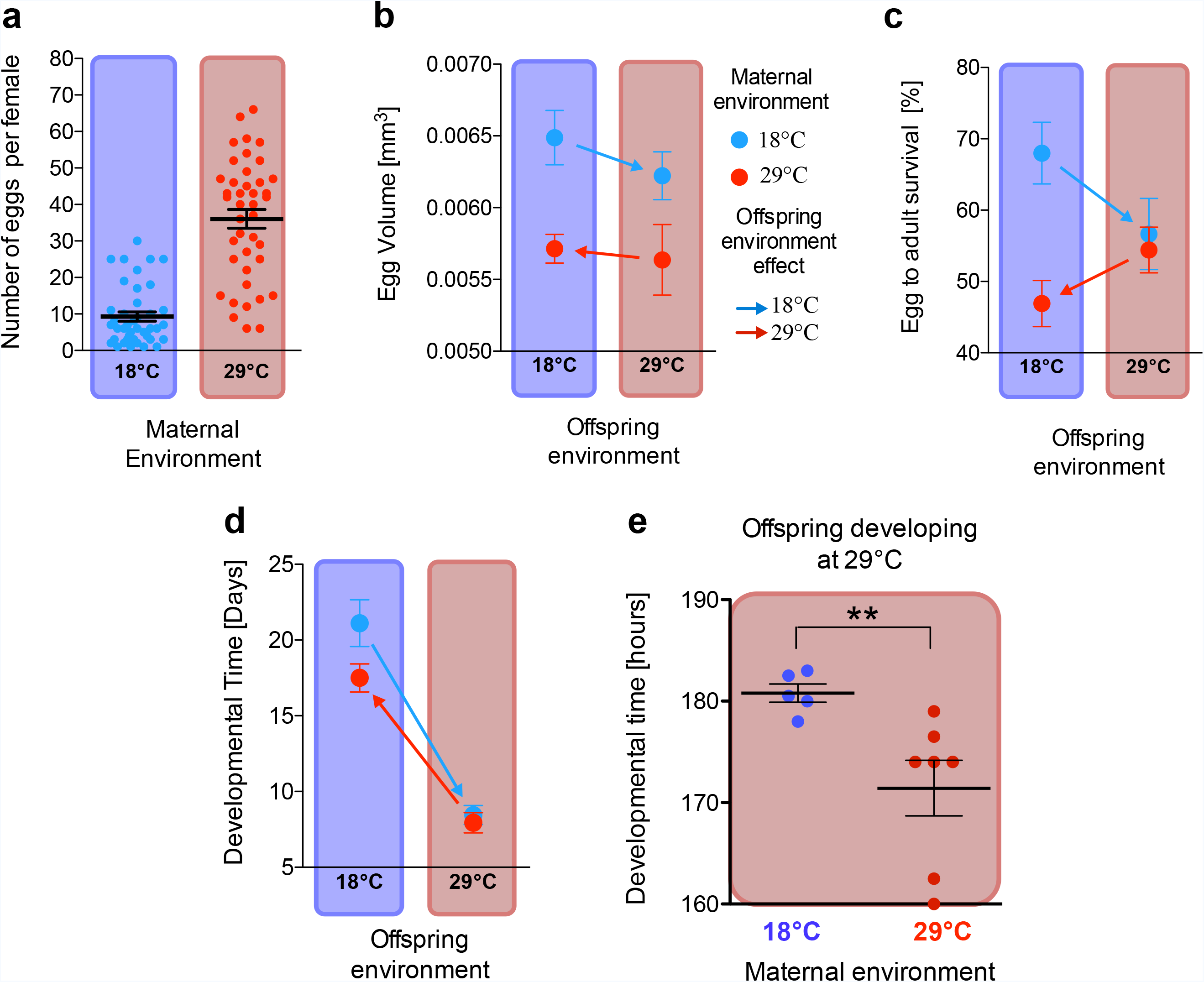
Influence of maternal temperature on egg phenotypes. *(A)* Average number of eggs laid in 24 hours by single females housed at 18°C or 29°C. Number of replicates is 41 females for each condition. *(B)* Effect of maternal and offspring conditions on egg volume. Mothers and eggs were housed at 18°C or 29°C as indicated. Arrows indicate direction of the change due to the mismatch of parents and offspring environments. The number of F_1_ mothers tested in each condition ranged from 11-31. Error bars indicate Standard Error of the Mean (S.E.M). *(C)* Effect of maternal and offspring conditions on offspring survival. The number of broods tested in each condition was 41. *(D)* Effect of maternal and offspring conditions on offspring developmental time. The number of clutches tested in each condition was 34. *(E)* Developmental time at 29°C of offspring from mothers housed at 18°C or 29°C. Each dot represents one egg. Mann-Whitney U-test indicates a significant effect of maternal condition on offspring developmental time (P=0.0089).

### Statistics

The unit of replication is the F_1_ mother. All graphs display the mean measure of offspring phenotypes per mother.

For statistical analysis, effects of treatments on the variables egg volume, progeny number (after Log-transformation), wing length and wing width were determined using a standard least square mixed effect model in which variables were continuous and normally distributed. Mother, offspring temperature conditions (18°C or 29°C) and offspring sex, as well as their interactions were modelled as fixed effects, and individual F_1_ mothers as random effects.

For survival (fig. 2*c*), a binomial logistic regression, with Mother and offspring temperature conditions as fixed effects and individual mothers as a random effect, was applied on the proportion of eggs that survived to adulthood.

Developmental time (fig. 2*d*) and grand offspring number (fig. 4) data showed unequal variance as determined by Bartlett test of homogeneity of variance. An Analysis of variance was performed on these data allowing for unequal variance using the Generalized Least Square function from the nlme package in R (R Studio Team 2016,v1.0.143). We used the varIdent variance function, which fits a separate residual variance for each of the four categories of the data. For testing significance of fixed effects, models were re-fitted with max likelihood and fixed effects were tested with Likelihood Ratio Test (LRT).

The variables offspring survival and developmental times were continuous and normally distributed. Differences between experimental conditions on these variables were determined using a standard least square model with mother and offspring temperature conditions (18°C or 29°C) modeled as fixed effects.

Unless indicated otherwise, Mixed Standard Least Squares models were run with JMP v. 9.0 for Mac, T-test and Mann-Whitney U-test were performed using GraphPad Prism (GraphPad software, Inc.). Effect sizes between two treatments were computed using

Cohen’s d formula: 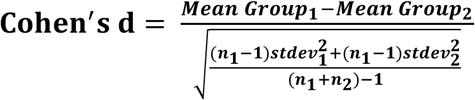

## Results

### Females lay fewer but larger eggs at 18°C than at 29°C

To determine the influence of temperature on reproduction of the F_1_ females we first analysed the number of eggs laid at 18°C and 29°C. As previously reported (Huey *et al.*, 1995), females laid significantly fewer eggs at 18°C than 29°C (Mann-Whitney test; U=127.5; p<0.0001)(fig. 2*a*). Eggs measured within 1 hour after laying had a larger volume when produced by mothers housed at 18°C than at 29°C (fig. 2*b*). To control for a direct early effect of temperature on egg volume independent of maternal effects, we placed eggs of both maternal temperatures in mismatched conditions for 5 hours (time between egg-laying and hatching is about 24 hours) directly after egg laying and compared their volume with that of matched eggs (fig. 2*b*). Maternal temperature condition had a significant effect on egg volume (fig. 2*b*; table 1), which was larger at 18°C than 29°C (fig. 2*b*). Statistical analysis yielded no effect of egg temperature condition indicating that eggs do not show intrinsic phenotypic plasticity in volume during the first 5 hours of development and that, as expected, egg size is solely under maternal control (fig. 2*b*; table 1).

**Table 1:** Test of between-subject fixed effects of maternal and offspring temperature conditions

| Phenotype | Factors<br>M=Mother condition<br>O=Offspring condition | D.F. | Test statistic | P |
| --- | --- | --- | --- | --- |
| Egg volume<br>(fig 1B) | *M* | 1,48.11 | F=33.56 | **<0.0001** |
|  | *O* | 1,48.11 | F=1.90 | 0.1738 |
|  | *M x O interaction* | 1,48.11 | F=0.34 | 0.5604 |
| Survivability<br>(fig 1C) | *M* | 1,3689 | Z=-3.825 | **0.01145** |
|  | *O* | 1,3689 | Z=-2.028 | **0.04259** |
|  | *M x O interaction* | 1, 689 | Z=2.839 | **0.00452** |
| Developmental<br>time<br>(fig 1D) | *M x O interaction* | 1,132 | LRT=51.404 | **<0.0001** |
| Wing length<br>(fig 2A&B) | *M* | 1,65.11 | F=23.40 | **<0.0001** |
|  | *O* | 1,58.36 | F=505.45 | **<0.0001** |
|  | *O sex* | 1,386.88 | F=281.43 | **<0.0001** |
|  | *M x O x O sex interaction* | 1,391.09 | F=6.80 | **0.0094** |
|  | *M x O interaction* | 1,65.11 | F=5.62 | **0.0206** |
| Wing Length<br>female<br>(fig 2A) | *M* | 1,61.17 | F=17.72 | **<0.0001** |
|  | *O* | 1,61.17 | F=292.23 | **<0.0001** |
|  | *M x O interaction* | 1,61.17 | F=0.61 | 0.4359 |
| Wing Length<br>male<br>(fig 2B) | *M* | 1,62.67 | F=17.72 | **<0.0001** |
|  | *O* | 1,62.67 | F=292.23 | **<0.0001** |
|  | *M x O interaction* | 1,62.67 | F=12.21 | **0.0009** |
| Wing width<br>Female<br>(fig 2C) | *M* | 1,55.91 | F=0.16 | 0,6828 |
|  | *O* | 1,55.91 | F=81.68 | **<0.0001** |
|  | *M x O interaction* | 1,55.91 | F=1.85 | 0.1784 |
| Wing width<br>male<br>(fig 2D) | *M* | 1,62.28 | F=0.02 | 0,8706 |
|  | *O* | 1,62.28 | F=169.74 | **<0.0001** |
|  | *M x O interaction* | 1,62.28 | F=0.41 | 0.5212 |
| Grand-<br>offspring<br>number<br>(Fig 3) | *M* | 1,93 | LRT=3.81 | 0.0507 |
|  | *O* | 1,93 | LRT=14.82 | **0.0018** |
|  | *M x O interaction* | 1,93 | LRT=0.17 | 0.6795 |

### Matched offspring have greater survival than mismatched ones

In matched conditions, survival is higher at 18°C than 29°C (Mann-Whitney test; U=537.5, P=0.0077) (fig. 2*c*), consistent with the documented deleterious effects of temperatures above 28°C (Petavy *et al.*, 2001). Mothers laying at 29°C might thus be making the best of a bad situation. More interestingly, there was a statistically significant interaction between maternal and offspring conditions on offspring survival indicating the presence of maternal effects in response to temperature (fig. 2*c*; table 1). These maternal effects suggest anticipatory matching because a mismatch between mother and offspring environments resulted in reduced offspring survival compared to matched conditions at both 18°C and 29°C (fig. 2*c*).

### Offspring and maternal condition interact in determining developmental time

Eggs developing at 29°C developed faster than those developing at 18°C, irrespectively of mothers condition, showing a strong direct effect of temperature on offspring development (fig. 2*d*; table 1; table S1). In addition, statistical analysis indicates a highly significant interaction between mother and offspring temperature conditions indicating maternal effects on offspring developmental speed, in addition to the direct effects of temperature on offspring development (fig. 2*d*; table 1). The developmental speed of offspring from mothers housed at 29°C, but who developed at mismatched 18°C, eclosed three days earlier than matched offspring from mothers housed at 18°C, whereas this was not the case for the 29°C developmental condition (fig. 2*d*).

The measurement of maternal effects on offspring developing at 29°C are less accurate that those at 18°C because of the greater speed of development. To verify maternal effects on the development time of eggs housed at 29°C, and to estimate these effects with greater accuracy, we collected eggs hourly and monitored development using 1hr time-lapse imaging. Mismatched offspring eclosed as adults 9 hours later than matched ones, confirming the presence of maternal effects at 29°C (fig. 2*e*).

Offspring temperature has the largest effect size on developmental speed, showing that intrinsic phenotypic plasticity is more important than maternal effects for this trait (fig. 2*d*; table 1; table S1). The maternal effect, however, did influence developmental speed, which is always faster in offspring from mothers housed at 29°C than offspring from mothers housed at 18°C, irrespectively of the temperature condition of the offspring themselves (fig. 2*d)*.

### Wing length but not width is influenced by maternal effects

Both wing length and size are significantly larger in individuals that developed at 18°C compared to those at 29°C (fig. 3; table 1), and females had significantly longer wings than males (fig. 3; table 1). There is therefore a strong influence of offspring temperature condition and sex on size. However the wing length of both females (fig. 3*a*) and males (fig. 3*b*) was also significantly influenced by maternal temperature conditions (table 1). The observation that female offspring from mothers housed at 29°C always had shorter wings than female offspring from mothers housed at 18°C indicates that maternal effects on female wing length might be carry-over effects from the temperature in which the mothers were housed. However maternal effects have a different effect on male offspring than female offspring as indicated by the statistical 3-way interaction between maternal and offspring conditions and sex on wing lengths as well as the post hoc test per sex indicating that in males, but not females, the mother and offspring condition interact to determine wing length (table 1). Male offspring from mothers housed at 18°C have larger wings than male offspring from mothers housed at 29°C, but only when the offspring was exposed to 29°C. Indeed, wing length does not significantly different between matched F_2_ males from mothers housed at 18°C or mismatched F_2_ males that grew at 18°C but that are from mothers housed at 29°C (t-test with Welch’s correction: t=1.303, df=79, P=0.196). The carry over effect from mothers housed at 29°C observed in females thus appears to be partly compensated in male offspring at 18°C.

**Figure 3:**
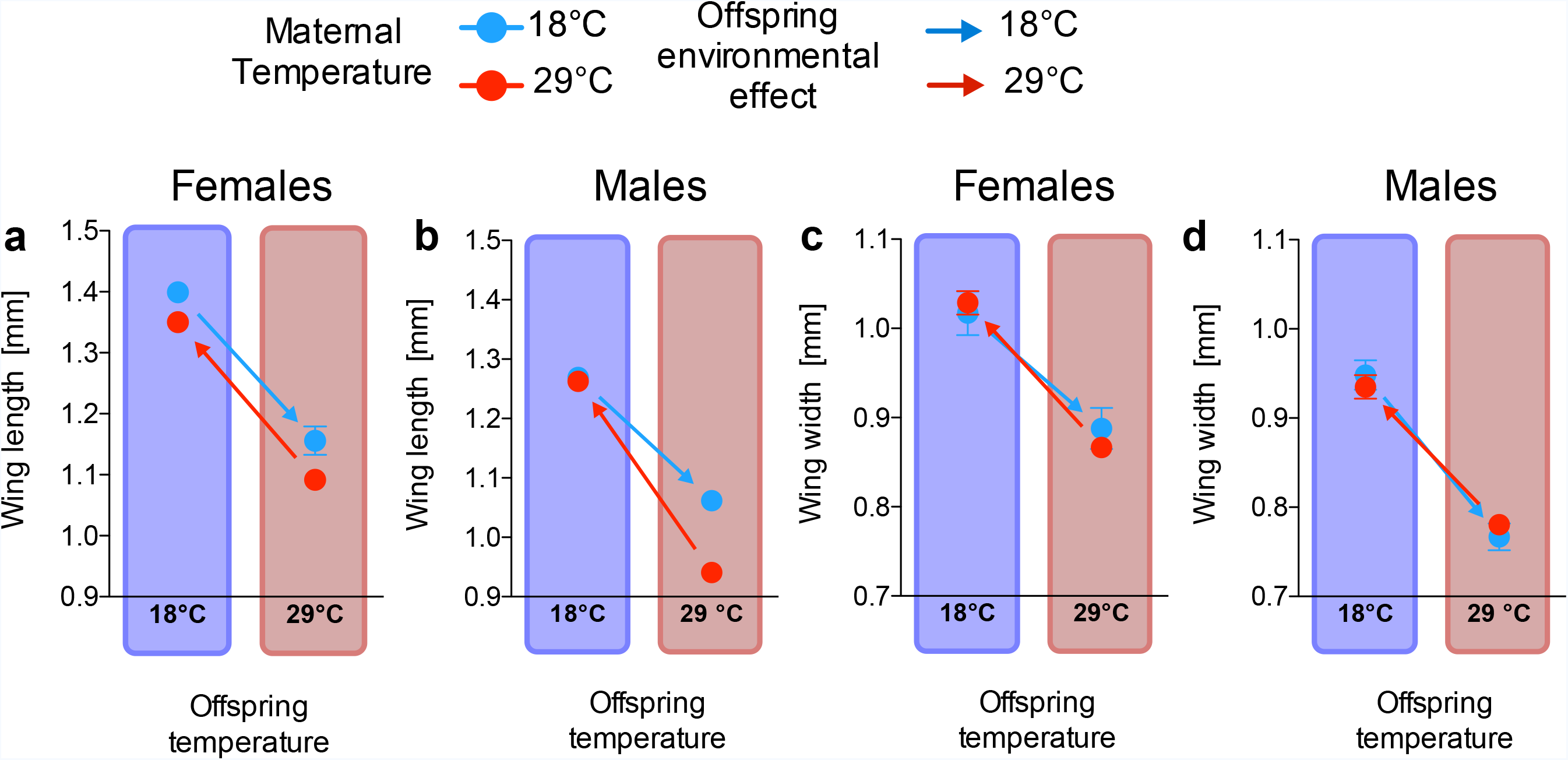
Influence of maternal and offspring temperatures on wing size. Mothers and eggs were housed in 18°C or 29°C environments as indicated. Arrows indicate direction of the change due to the mismatch of parents and offspring environments h. Error bars indicate Standard Error of the Mean (S.E.M). The number of replicate mothers was 16 in all 4 mother-offspring temperature combinations in Panels *(A-D)*.

There is no statistical effect of mother condition on wing width, neither by itself or in interaction with offspring condition (fig. 2*c*-*d*; table 1), but a strong effect of offspring condition alone indicating that individual differences due to temperature conditions are the result of intrinsic offspring phenotypic plasticity.

### Reproductive performance of F_2_ offspring is unaffected by F_1_ maternal condition

We determined the fecundity of matched and mismatched F_2_ offspring in the context of assortative sibling mating (fig. 4). Statistical analysis indicated a significant effect of F_2_ rearing condition but no effect of F_1_ mother condition (table 1). Within temperature conditions, matched and mismatched F_2_ offspring did not differ significantly in offspring number indicating a lack of F_1_ maternal effect extending to the F_2_ generation (fig. 4). Intriguingly, both matched and mismatched F_2_ offspring produced slightly more F_3_ offspring at 29°C than at 18°C (fig. 4), suggestive of decreased fecundity at 18°C as a result of intrinsic phenotypic plasticity.

**Figure 4:**
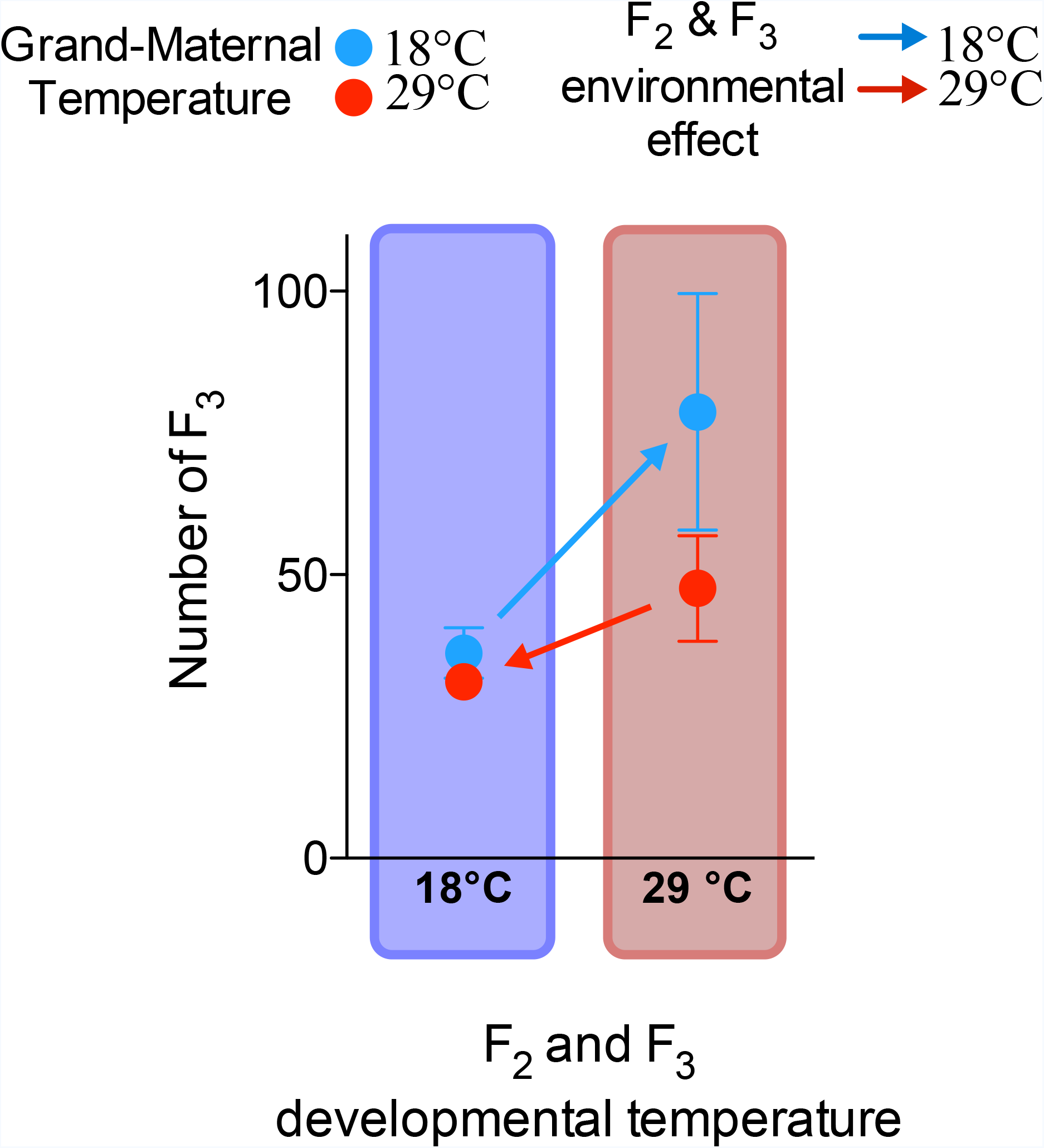
Reproductive performance of F_2_ offspring. Single Matched and mismatched F_2_ females were mated singly with their brother and led in the conditions in which they developed. Females laid their eggs and the eggs developed in the same conditions. The number of adult offspring was counted at emergence. Error bars indicate Standard Error of the Mean (S.E.M). The number of replicate F_1_ mothers ranged from 19 to 32.

## Discussion

The goal of the present study was to test, in a laboratory setting, the extent to which anticipatory maternal effects in *Drosophila melanogaster* may modulate phenotypic values in their offspring traits in response to temperature - an environmental variable known to have relevance for fitness (Kingsolver & Huey, 2008). We used a full experimental match-mismatch design allowing us to separate maternal effects from intrinsic offspring plasticity and maternal adjustment from carry over effects. Evidence for matching, also known as anticipatory maternal effect, would come from mothers modifying offspring traits such that offspring reared and living in the same environment as that of their parents will have higher fitness than offspring living in an environment different from that of their parents (Mousseau and Dingle 1991; Leroi et al. 1994; Huey et al. 1999). We found that survival from egg to adult is subjected to anticipatory maternal matching in that offspring raised in the same temperature as their parents had a higher survival than those raised at different temperatures, irrespective of the actual temperature. Evidence for anticipatory effects was however not found for other phenotypes such as adult body size or fecundity. This latter is in keeping with previous work in *Drosophila*, which studied the consequences of parental effects in response to temperature on several phenotypic traits (Crill *et al.*, 1996) and on fitness (Gilchrist & Huey, 2001) and found evidence against adaptive matching but in favour for a higher fitness of flies whose parents were in hot conditions. These studies, however, measured fitness in terms of per capita rate of population increase but did not measure survival from egg to adult as we did.

The relative larger egg volume of mothers housed at 18°C compared to mothers housed at 29°C indicates that females provision eggs more at 18°C than at 29°C (fig. 2*b*). The effect size of temperature on egg volume and number are similar but in opposite directions suggesting the trade-off between egg volume and number found in other egg laying species (Williams, 2001)(table S1). This differential provisioning may provide maternal input to the offspring affecting developmental plasticity. Egg volume increases in response to selection for fast development in *Drosophila* (Bakker 1969) and a larger volume has a positive effects on embryonic viability and development rate, hatchling weight, larval feeding rate, and larval and pre-adult development rates (Azevedo *et al.*, 2010). This association between larger egg volume and higher survival is observed in our experiments where the smaller eggs produced by mothers at 29°C have lower survival to adulthood than those produced by mother housed at 18°C (fig. 2*c*). The low egg to adult survival at 29°C in our study is in keeping with previous reports of lower viability in conditions above 28°C (Petavy *et al.*, 2001). Another possible explanation for the differential survival at the different temperatures are differences in egg density due to lower egg-laying at 18°C than at 29°C; too many larvae can affect viability through food limitation (Horváth & Kalinka, 2016). The mean number of eggs per vial was lower (~8) at 18°C than at 29°C (~20), but corresponded to egg density that are far from leading to food limitation (starting at 175 eggs/vial) (Horváth & Kalinka, 2016). The match-mismatch design indicates the presence of anticipatory maternal effects because within one temperature condition, offspring raised in conditions that match that of their parents are more likely to survive development than those that are mismatched. This parental effect on survival is substantial and larger than the direct effect size of temperature on offspring survival, indicating the relevance of parental effect in offspring adaptation to temperature (table S1). As survival is a close proxy for fitness, it suggests that anticipatory parental effects can participate to evolutionary adaptation.

Maternal condition had a significant effect on developmental speed indicative of carry-over effects because both matched and mismatched offspring from mothers housed at 18°C developed slower than both matched and mismatched offspring from mothers housed at 29°C (fig. 2*d-e*). Mothers housed at 18°C thus slow down offspring development and mothers housed at 29°C speed it up. Our combined data on developmental speed and survival (fig. 2*c-e*) may however suggest anticipatory maternal effects on offspring development. Intrinsic offspring phenotypic plasticity has a larger effect on developmental speed than maternal effects (fig. 2*d*; table S1), but anticipatory maternal effects have a large effect on survival compared to intrinsic phenotypic plasticity (fig. 2c; table S1). Reduced survival when the offspring environment is mismatched with that of the mother (fig. 2*c*) might therefore stem from maternal effects interfering to slow down development in, for instance, the anticipated colder conditions, increasing viability, while the hotter temperature in which the offspring is actually developing directly increases offspring development speed (and vice versa for mismatched offspring from mothers housed at hotter temperatures)(fig. 2*d-e*). Incompatibility between these two processes might be the cause of the decreased survival when maternal and offspring environments are mismatched. Anticipatory maternal matching might be a normal feature of *Drosophila* development and the basis for the greater survival of offspring developing in conditions matched with those of their parents (fig. 2*c*). This might be an adaptation to the ecological conditions in which *Drosophila melanogaster* lives, which involves feeding and developing on fermenting food substrates where a fast development is crucial to outcompete microbes and fungi (whose growth is also influenced by temperature)(Markow & O’Grady, 2008). Mothers may be able to prime their eggs for a faster or slower rate of development, through a mechanism that affects egg volume, that can be predicted from temperature conditions at the time of egg production, which would trigger a cascade of adaptation to higher temperatures in the larvae such as changes in feeding and developmental rate (Azevedo *et al.*, 2010).

Adaptive matching has a large effect on early viability, but do these effects persist well into the adult stage? One of the largest effects of temperature on adult size is that flies are bigger when the developmental conditions are cooler (Kingsolver & Huey, 2008
). This is confirmed in our experiments showing that the major effect on adult wing size comes from offspring temperature conditions (fig. 4; table 1; Table S1). Maternal condition also had an effect on offspring adult wing size, albeit smaller (fig. 4; table S1). Given the high mortality observed at 29°C (fig. 2c), the observation of smaller wings at this temperature could have been the result of temperature selecting for flies with smaller wings, instead of a result of phenotypic plasticity. This is however unlikely to be the case because we used a wild-type strains that is largely inbred, thus reducing the difficulty in separating parental effects from selection on offspring genotype during the experiments (Faurby *et al.*, 2005). Maternal effects can be expected to influence adult offspring phenotype because final adult size is regulated by the size at which the larva stops growing and initiates metamorphosis. As the decision to metamorphose is made earlier in the final instar larva (Mirth & Shingleton, 2012), maternal effect on egg composition could still be acting on growth. However, this effect is not anticipatory matching but rather a carry-over effect because female offspring from mothers housed at 18°C always have longer wings than offspring from mothers housed at 29°C (fig. 4*a-b*). The carry over effect appears buffered in male offspring, since males that developed at 18°C had similar wing lengths whether they originated from a mother housed at 18°C or 29°C (fig. 4*a-b*). Males buffering carry-over maternal effects on wing length might give them an advantage because male-male competition and female mate choice is influenced by male wing and body size (Roff, 1986). However, males from mothers housed at 29°C and developing at 29°C have smaller wing size than those from mothers housed at 18°C. Wing area and length contribute to adaptation to temperature conditions because larger wings improve flight performance at colder conditions (Frazier *et al.*, 2008). Males with larger body size (and wing) have higher mate competitive advantage (Kingsolver & Huey, 2008), which may select for mothers influencing their sons to have the greatest possible wing size for the perceived temperature. A male developing at 29°C, might have still be primed by his mother to develop greater wing size.

Given the observation of anticipatory maternal effects on temperature conditions, one outstanding question remains their potential fitness significance. By measuring the number of F_2_ and F_3_ offspring produced in different temperature conditions and under match or mismatched condition, we can determine the relative fitness consequences of maternal effects in matched and mismatched conditions. In matched conditions, the 29°C temperature leads to 3 times more F_2_ offspring than 18°C, leading to the clear conclusion that hotter temperature is conducive to higher fitness (fig. 5). In mismatched conditions, mothers housed at 29°C also have more offspring than mothers housed at 18°C confirming previous observations that parents under hotter temperatures will have more offspring irrespective of offspring conditions (Gilchrist & Huey, 2001; Marshall & Sinclair, 2010)(fig. 5). However, within offspring condition, comparison of F_2_ production of matched vs mismatched offspring always shows an advantage for matched offspring resulting in 1.2 times increases in progeny (fig. 5). This indicates that matching the temperature conditions of parents and offspring has fitness benefits for the parents, supporting the adaptive matching hypothesis. But does adaptive matching have an effect on the offspring fitness (F_2_)? This can be derived from comparing the number of offspring (F_3_) from F_2_ parents raised in matched 18°C vs mismatched 29°C conditions, since the only difference between these two treatments is the condition of the F_1_ mother. In this case, matched F_2_ parents have slightly more offspring than mismatched F_2_ parents (1.2 times more; fig. 4), indicating potential transgenerational fitness benefits of matching. However this effect is not statistically significant (as already determined in fig. 4; table 1). Comparing the number of offspring (F_3_) from F_2_ parents raised in matched 29°C vs mismatched 18°C conditions shows that matched 29°C F_2_ parents had fewer offspring (0.6x) than mismatched one (fig. 5), arguing against the adaptive matching hypothesis. However the effect of F_1_ mother is again not statistically significant, indicating of a lack of negative maternal influence. We therefore conclude that this is an indication that there are little to no fitness consequences of adaptive matching on the offspring, just on the parents. The short generation time of *Drosophila* and the natural fluctuation in temperature conditions might make maternal effects efficient for short term adaptation to developmental conditions of the offspring but not for its reproductive ability as an adult. Committing those effects to the next generation might be futile given that the conditions are likely to have changed again. A test of the adaptive value of these anticipatory effect will be to demonstrate that the population used has been subject to natural selection in a variable, but predictable, environment. As we used an inbred fly strains that has been kept in the lab for a long time, we cannot reach this conclusion.

**Figure 5:**
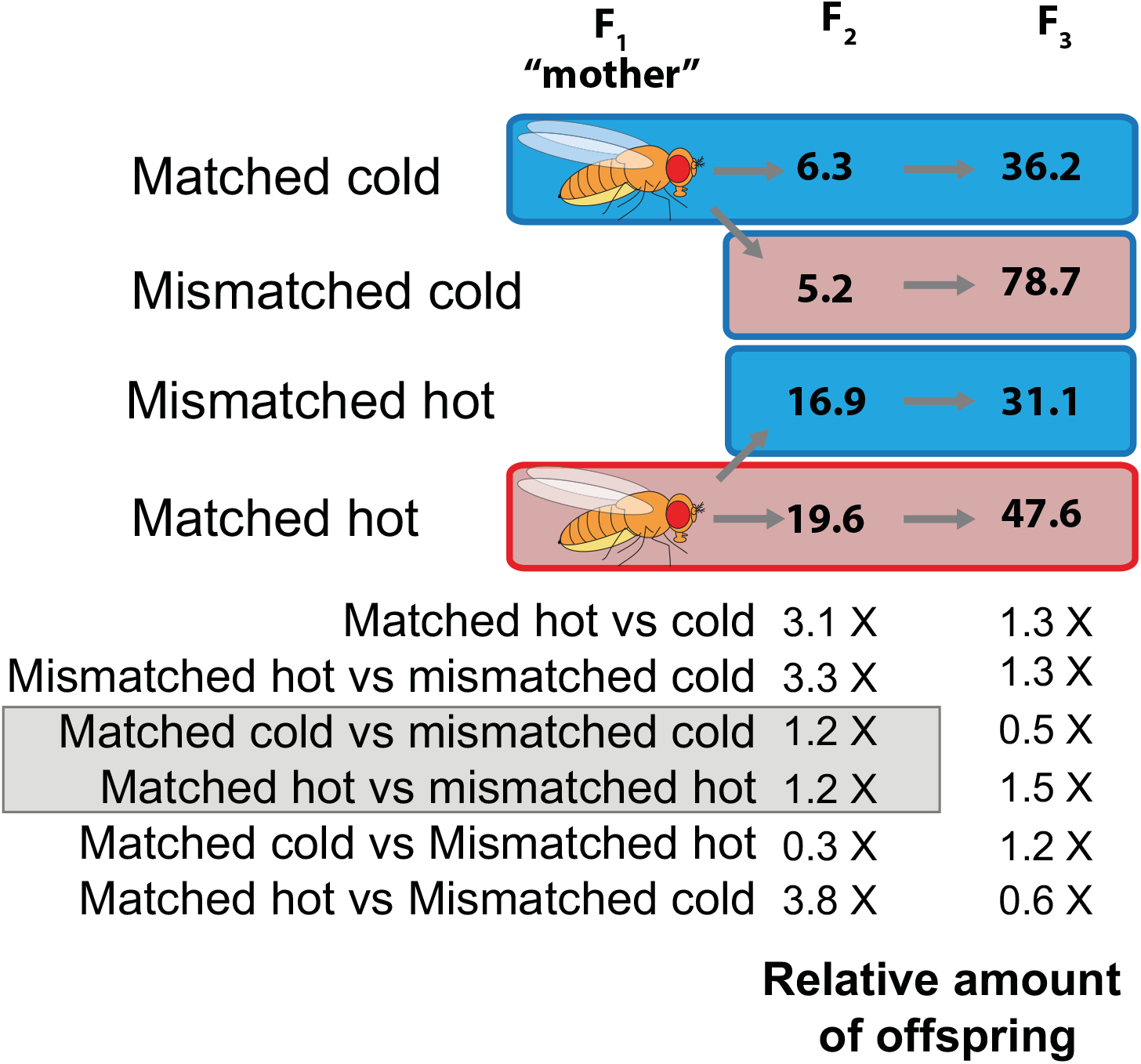
Fitness consequences of maternal effects. The temperature condition of the mother is indicated by the border colour (Blue for 18°C and red for 29°C). The colour of the boxes themselves indicates the condition in which the offspring developed and reproduced (Blue for 18°C and red for 29°C). A difference in colour between borders and shading indicates a mismatch condition. Numbers in the boxes in the first two columns indicate the number of offspring produced by a single F_1_ or F_2_ mother. Below the graph are relative differences in offspring production between the different treatments discussed in the text. The grey box highlights treatments whose comparison reveal maternal effects.

In summary, our results suggest the existence of anticipatory maternal effects in response to temperature in *Drosophila melanogaster.* These maternal effects affect mostly parental fitness, by increasing offspring survival without increasing offspring fecundity. Adaptive matching parental effects to temperature are thus not multigenerational. We could only find anticipatory matching in the context of survival but suspect that maternal effects on developmental speed, that may appear as carry over effects, might be connected to an early maternal effect that sets embryos in a developmental trajectory that is adapted to the temperature conditions experienced by the mother. A better mechanistic understanding of maternal effects is therefore required to distinguish between anticipatory and carry-over effects. Given the breadth of mechanistic knowledge on the effects of the maternal genome on early *Drosophila* development and the tools available to study *Drosophila* development (Schüpbach & Wieschaus, 1986), a mechanistic understanding of anticipatory maternal effects should now be on the horizon. It will be equally relevant to demonstrate the adaptive significance of these effects observed under laboratory conditions by showing, in an outbred population, that anticipatory maternal effects can be selected in environments that are variable, but predictable.

## Acknowledgement

We thank Pinar Güler, Martine Maan, Bernd Riedstra for critical comments on the manuscript, and Ido Pen for help with statistics. This work was supported in parts by an Erasmus Mundus Svagata short-term post-doctoral fellowship to Snigdha Mohan, as well as funds from the University of Groningen and the Dutch organization for scientific research (NWO) to J.C. Billeter.

## Author contributions

A.G.G. and J.-C.B designed and interpreted the study. S.M. and C.V. performed all experiments. A.G.G. and J.-C.B performed the statistical analysis. J.-C.B. prepared the figures and manuscript.

